# Eye-hand coordination beyond vision: preceding gaze history shapes reaching memories

**DOI:** 10.1101/2025.11.11.687771

**Authors:** Naotoshi Abekawa, Hiroaki Gomi

## Abstract

Skilled reaching behaviors, such as grasping a cup or catching a ball, require adaptive changes in response to error signals. In natural behavior, reaches are typically coordinated with preceding eye movements toward the target. Despite this behavioral relevance, the potential linkage between preceding eye movements themselves, rather than static gaze positions, and memory formation for reaching has been largely overlooked. Here, we used a visuomotor rotation task to investigate how recent dynamic gaze states contribute to the formation and retrieval of reaching memories. We show that, in the learning of two conflicting visuomotor maps that would normally interfere with one another (namely, clockwise and counterclockwise rotation), not only the static spatial relationship between the gaze and the target but also the dynamic state of the eye movements themselves can serve as contextual cues that enable learning. We further show that the generalization pattern of reach learning across different gaze conditions can be explained by the gaze state’s distance computed with temporal weighting. These findings indicate that reaching memories are encoded in tight association with the recent history of preceding gaze states. Our results expand the functional role of eye-hand coordination beyond visual guidance, revealing its importance in orchestrating flexible motor learning.

**Significant Statement:** In everyday behavior, goal-directed arm movements are typically preceded by eye movements, and eye-hand coordination has been regarded mainly as a functional mechanism for visual guidance of action. This study expands that view by showing that preceding gaze states also shape how the brain learns and retrieves motor memories of reaches. We demonstrate that conflicting visuomotor maps, which are usually difficult to learn because of memory interference, can be learned simultaneously when they are associated with different preceding eye movements. Furthermore, generalization experiments show that reach learning depends on the recent history of gaze states within a brief time window before reach onset. These findings indicate that preceding gaze history plays a significant role in shaping flexible reach memories.

## Introduction

In an ever-changing world, skillful arm-reaching movements rely on motor learning, which involves the formation and updating of sensorimotor maps called internal models (Kawato, 1999). A key issue in motor learning is understanding how the brain represents internal models, particularly their association with various body states (Shadmehr, 2004; Krakauer et al., 2019). Reaches are typically preceded by eye movements toward the target, hence much research has examined how the brain spatiotemporally coordinates eye and hand movements, both in daily activities and laboratory tasks, particularly in relation to target representation (Land, 2009; de Brouwer et al., 2021). Similarly, reach learning often occurs under coordinated eye-then-reach movements (Davids et al., 2005). For example, tennis players track a moving ball with their eyes while learning and executing skilled reaching actions. This natural eye-hand coupling raises an unexplored question that spans both eye–hand coordination and motor learning: Does the preceding dynamic gaze state contribute to the encoding of motor memories for subsequent reaching movements?

There are two opposing views on this issue. One emphasizes the role of gaze in target representation and deems it unrelated to motor learning. From this view, gaze provides target localization and a reference frame for encoding target location. Target locations are initially represented in a gaze-centered frame and updated through eye movements (Crawford et al., 2004, 2011; Andersen and Cui, 2009). Such gaze-dependent signals are transformed into gaze-independent signals (e.g., hand-to-target trajectories), subsequently used with internal models to generate reach motor commands. In this framework, preceding gaze states are not expected to contribute directly to motor learning.

An alternative view focuses on the coupling of sensorimotor signals across eye and hand motor systems. Such motor-coordination mechanisms enable multiple effectors to work cooperatively, as observed in bimanual and multi-joint movements (Diedrichsen et al., 2010). From this view, motor commands to one effector depend on the state of another. Thus, in tightly coupled eye-hand coordination (Miall et al., 2001; Ariff et al., 2002; Yttri et al., 2013), preceding gaze states may play a role in representing reach internal models.

Consistent with the latter view, motor-learning studies have demonstrated that learning reaching movements with one arm depends on the state of the other arm (Nozaki et al., 2006; Yokoi et al., 2011; Gippert et al., 2023; Howard et al., 2024). In context-dependent learning paradigms used in these studies, two opposing perturbations (i.e., clockwise and counterclockwise velocity-dependent force fields) that normally interfere with each other can be learned simultaneously when each is associated with a distinct movement pattern of the other arm. Importantly, contextual movements can facilitate learning even when they occur before the adapted movement (Wainscott et al., 2005; Howard et al., 2012, 2024; Gippert et al., 2023).

These findings suggest that prior hand states facilitate the separation of motor memories and thus form part of internal-model representations. This is consistent with recent theories that emphasize preparatory neural states in motor-related cortices in motor learning (Sheahan et al., 2016; Vyas et al., 2020; Sun et al., 2022). Most studies, however, have focused on hand-related states as the learning context, and it remains untested whether similar parallel conclusions can be drawn for other body states. Although we recently demonstrated that the gaze location relative to the target (i.e., reaching towards a central or peripheral target) can separate visuomotor memories, the investigation was confined to specific static gaze conditions (Abekawa et al., 2022). Thus, it remains unknown whether/how preceding dynamic gaze states contribute to reach learning.

Here we demonstrate through a series of experiments that participants can learn two opposing visuomotor maps when each is associated with a distinct preceding dynamic gaze state. This eye-dependent learning effect is sensitive to the time interval between eye and reaching movements. Our findings provide evidence that memory encoding for reaching is tightly linked to the recent history of gaze states. Preliminary findings from Experiment 1 were previously reported in a conference proceeding (Abekawa and Gomi, 2024).

## Materials and Methods

### Participants

A total of 84 healthy right-handed individuals participated in Experiment 1 (including three experimental groups), Experiment 2, Experiment 3, Control Experiment 1 and Control Experiment 2, each with 12 participants. The sample size was chosen on the basis of our previous similar experiments and the current counterbalanced experimental design. All participants were naïve to the purpose of the study. They gave informed consent to participate, which was approved by the NTT Communication Science Laboratories Research Ethics Committee.

### Apparatus

Participants moved a stylus with their right hand on a digitizing tablet (Wacom Intuos 2) placed horizontally on the table (Figure 1A). The position of the stylus captured at 100 Hz was presented as a hand cursor on a vertical CRT monitor (Mitsubishi RDF223G; 100 Hz vertical refresh rate) placed approximately 50 cm from participants’ eyes. Participants were asked to move the cursor (black circle with 0.2-cm radius) from a start point (circle with 0.45-cm radius) to a target (white circle with 0.5-cm radius), where the cursor was moved upward or rightward by the forward or rightward stylus motion correspondingly. The hand and stylus were not directly visible. Participants’ heads were stabilized by a chin rest on which a video-based eye tracker (Eyelink 2000) was mounted to record gaze position (500 Hz). According to the trial type, participants were asked to keep fixating on a fixation cross (0.8-cm x 0.8-cm) or to make saccades. Stimulus presentation and experimental procedure were controlled using MATLAB (R2012a, MathWorks) and Psychophysics Toolbox (PTB-3)

**Figure 1.**
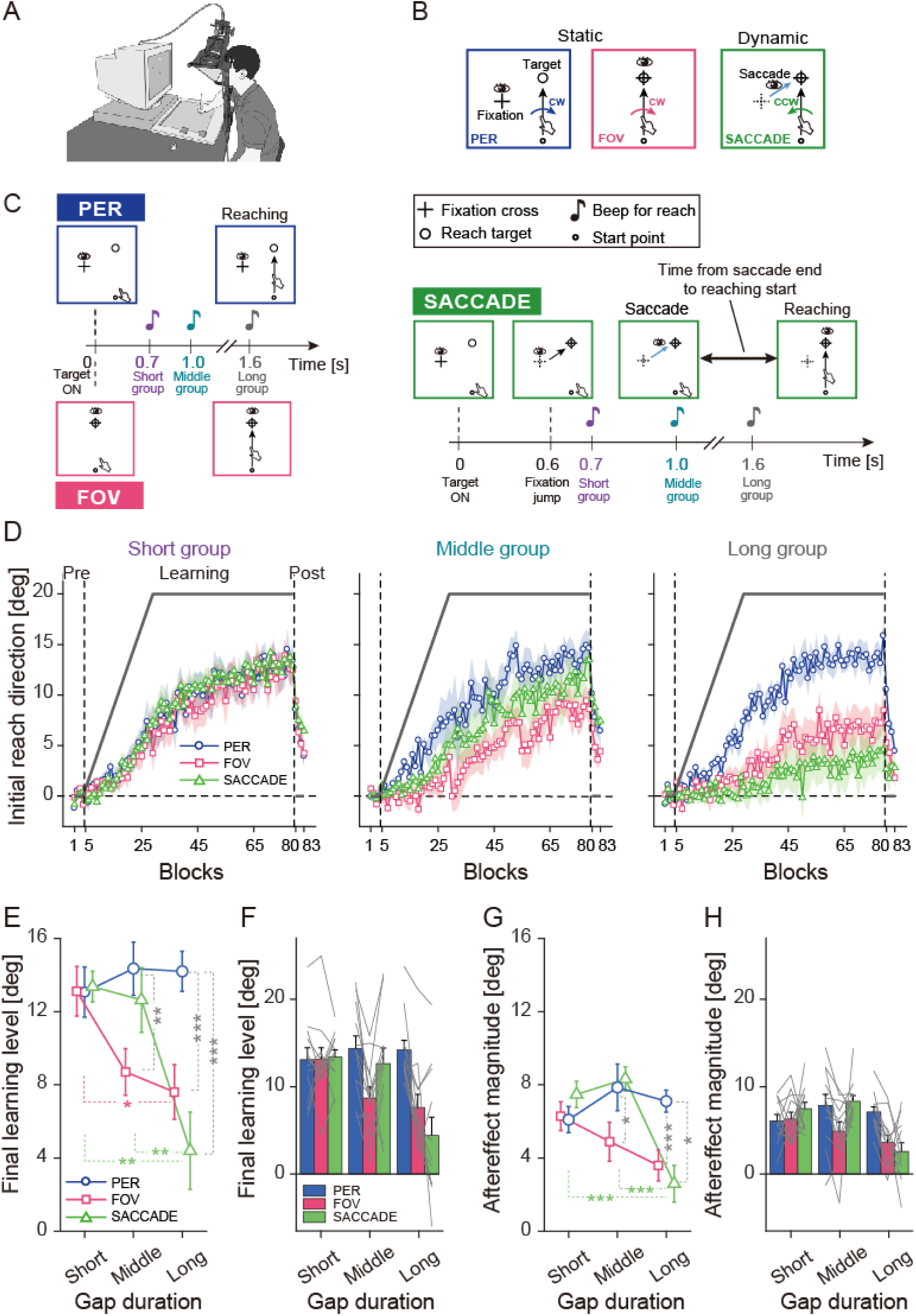
Experiment 1: Simultaneous learning based on static and dynamic gaze states. **A,** Experimental apparatus. Participants performed reaching movements on a digitizing tablet to move a visual cursor toward a target on a monitor. **B,** Gaze conditions. Under static conditions, reaching was performed with gaze fixed on the target (FOV) or elsewhere (PER). Under the dynamic condition (SACCADE), participants made a saccade toward the reach target then reached toward the foveal target. Three gaze conditions were interleaved randomly across trials. Visuomotor rotations were applied, with their directions associated with each condition (curved arrows; CW for static and CCW for dynamic; See main text for details). **C,** Trial sequence. In PER trials, the trial started when participants maintained fixation and the reach target appeared. Reaching was then initiated following a go beep. FOV trials were identical to the PER trials except that fixation was on the reach target. In SACCADE trials, participants made a saccade in response to a fixation jump then reached toward the target at the go beep. The go beep timing depended on the gap-duration group (Short, Middle, or Long) to manipulate the gap duration between saccade end and reach initiation. Each participant was assigned to one group. **D,** Learning curves for each gaze condition across three gap-duration groups. Initial reach direction was used as the learning measure, with its sign adjusted to match the rotation direction (positive values indicate learning). Data were averaged within each block and plotted as mean ± SEM across participants. The angle of visuomotor rotation gradually increased up to 20° (black line). **E,** Final learning levels of initial reach direction (averaged over the last four blocks in the learning session), shown for a two-way mixed ANOVA with gaze condition (PER, FOV, and SACCADE; within-participants) and Gap duration (Short, Middle, and Long; between-participants). Data are presented as mean ± SEM across participants. Asterisks indicate significant differences (Tukey post hoc tests; * p<0.05, ** p<0.01, *** p<0.001). **F,** Final learning levels (same data as in E) replotted for each Gap duration, with individual participant data shown as gray lines. **G, H** Aftereffect magnitudes (averaged over the four blocks of the post-learning session), shown in the same format as in E and F.

### Experimental design

#### Experiment 1

Experiment 1 was designed to investigate whether two opposing visuomotor rotations can be learned when each was associated with dynamic and static gaze context. A total of 36 participants were assigned to one of three timing groups (12 individuals per group): Short (8 females; age 33.5 ± 10.3, mean ± SD), Middle (6 females; age 30.2 ± 11.2, mean ± SD), and Long (8 females; age 35.9 ± 11.6, mean ± SD). The timing of the “go” cue to initiate the reach varied across groups to manipulate the relative timing of eye and hand movements. Details on this timing group are described later.

In each trial, participants performed the reaching task under one of three different gaze conditions (Figure 1B). Under the two static conditions (PER and FOV), participants reached toward the target while maintaining gaze fixation at each fixation cross throughout the trial. Under the dynamic condition (SACCADE), they made a saccadic eye movement to the target, followed by reaching toward the foveal target. The reach target was always located 10 cm above the start point on the screen. For half the participants, the fixation cross under PER and SACCADE was presented on the left side of the target, 10 cm from the start point (i.e., 7.66 cm left and 3.57 cm below the reach target), as shown in Figure 1B. Participants reached toward the target in the right visual field under PER, and under SACCADE they first made a rightward saccade, followed immediately by a reach toward the foveal target. To rule out the potential confounding effect of the fixation position under PER and saccade direction under SACCADE, the location of initial fixation was vertically mirrored (i.e., lower right to the reach target) for the other half of participants.

Each trial started with participants placing the cursor at the start point and directing the gaze to the fixation cross (Figure 1C). After maintaining fixation for 0.7 s, a reach target appeared. Under the static conditions (i.e., PER and FOV; Figure 1C, left), after a delay according to the timing group, a beeping sound was given for initiating the reach. Thus, participants reached toward the peripheral target under PER and foveal target under FOV and terminated the reach at a second beep presented 0.3 s after reach onset. Under the dynamic condition (i.e., SACCADE; Figure 1C, right), the trial started in the same way as under PER, but the fixation cross jumped to the target 0.6 s after the target presentation. Participants had to make a saccade to the new fixation position (i.e., reach target) as fast as possible and initiate the reach in response to the subsequent beep. In this time course, the reach began after saccade termination; thus, under SACCADE, participants first made a saccade then performed the reach toward the foveal target.

Throughout the experiment, the three gaze conditions (PER, FOV, and SACCADE) were randomly presented within each block of 16 trials, consisting of 4 PER, 4 FOV, and 8 SACCADE trials. The experiment comprised three sessions. After familiarization, participants completed a pre-learning session (4 blocks/64 trials) with normal visual feedback. In the subsequent learning session (75 blocks/1200 trials), a visuomotor rotation was applied to the cursor movements. The rotation angle increased by 0.8° per block over 25 blocks to reach 20° then was maintained for the remaining 50 blocks. The rotation direction (CW or CCW) varied pseudo-randomly across trials but was uniquely associated with the gaze conditions. For half the participants (n=6), CW was applied for the static gaze conditions (i.e., PER and FOV) and CCW for the dynamic gaze condition (i.e., SACCADE), and this association was reversed for the other half. To ensure an equal number of CW and CCW trials in the simultaneous learning paradigm, the number of trials for each gaze condition within a block (4 PER, 4 FOV, and 8 SACCADE) was set. Finally, participants completed a post-learning session (4 blocks/64 trials) identical to the pre-learning session (i.e., normal feedback).

The rationale for including these three conditions, with CW rotations assigned to the two static gaze conditions (PER and FOV) and CCW rotations assigned to the dynamic gaze condition (SACCADE), was to dissociate the contribution of the dynamic gaze state from that of the static gaze states. Our previous study demonstrated that distinct static gaze states (i.e., PER and FOV) are sufficient to support simultaneous learning of opposing visuomotor rotations (Abekawa et al., 2022). A saccade to the target is necessarily accompanied by two static gaze states: fixation of the peripheral target before the saccade and fixation of the foveal target after the saccade. Thus, SACCADE in our paradigm inevitably contains three potential contextual gaze-state cues: the pre-saccadic static peripheral fixation state, the dynamic saccadic state, and the post-saccadic static foveal fixation state. Therefore, if only two gaze conditions were used as a paradigm (e.g., SACCADE with CW and PER with CCW), even if dual learning were observed, it would remain unclear whether learning was supported by the contextual difference between the static gaze state in PER and the dynamic gaze state in SACCADE, or simply by the difference between the static gaze states in the two conditions (i.e., peripheral fixation in PER versus post-saccadic foveal fixation in SACCADE). Likewise, using only SACCADE and FOV would not exclude the possibility that the pre-saccadic static fixation state on the peripheral target, which is also included in SACCADE, served as the contextual cue. Therefore, simply testing learnability by pairing SACCADE with either one of the static fixation conditions would not provide evidence that the dynamic gaze state itself can serve as a contextual cue for visuomotor rotation learning. In contrast, in the abovementioned three-condition design, if a visuomotor map specific to SACCADE is learned separately from those associated with the two static fixation conditions, this would indicate that participants used the dynamic gaze state as a contextual cue for learning.

We further hypothesized that the contextual learning effect in SACCADE would depend on the interval between saccade termination and reach initiation. As this interval increases, the contextual information provided by the dynamic gaze state may decay over time, leaving the post-saccadic foveal fixation state as the dominant contextual cue. Consequently, when this interval is sufficiently long, the SACCADE context becomes increasingly similar to the FOV context, leading to interference between the two and reduced learning. In contrast, learning in PER would remain unaffected because reaching to a peripheral target provides a distinct contextual cue relative to the other two gaze conditions, as reported in previous work (Abekawa et al., 2022). To test this hypothesis, participants were assigned to Short, Middle, and Long gap groups, in which the “Go” beep for reaches was presented 0.7, 1.0, and 1.6 s after the target presentation, respectively (Figure 1C, right). The resulting interval between saccade termination and reach onset was 264 ± 97 ms (91–408 ms) for the Short, 541 ± 49 ms (466–638 ms) for the Middle, and 1213 ± 95 ms (109 –1449 ms) for the Long (mean ± SD and range across participants). To match the time course across conditions, the timing of the beep also varied according to the three timing groups under PER and FOV, which did not require saccades (Figure 1C, left).

Trials that violated any of the following criteria were specified as errors with a warning beep and repeated at the end of the block: 1) reach movement time was less than 0.15 s or more than 0.7 s, 2) the reach reaction time was over 1 s, 3) saccade reaction time was over 0.4 s, 4) eye movements were detected under PER and FOV, and 5) reach initiated before saccade termination under SACCADE.

#### Control Experiment 1

Control Experiment 1 was conducted to rule out the possibility that the visual cues of fixation cross jumps, rather than eye movements, served as a contextual effect in Experiment 1. Twelve participants (9 females; age 33.4 ± 9.4, mean ± SD) took part in this experiment. The task was identical to that of the Short group in Experiment 1, except that SACCADE was replaced with FIX-JMUP (Figure 2A). Thus, there were three gaze conditions: PER, FOV, and FIX-JUMP. Under FIX-JUMP, the trial began as under PER, but 0.6 s after target presentation the fixation cross jumped to the target. Participants had to maintain gaze fixation at the original location of the cross prior to the jump. The direction of visuomotor rotation was associated with static and visual dynamic conditions (i.e., CW for PER and FOV, CCW for FIX-JUMP), with the assignment counterbalanced across participants.

**Figure 2.**
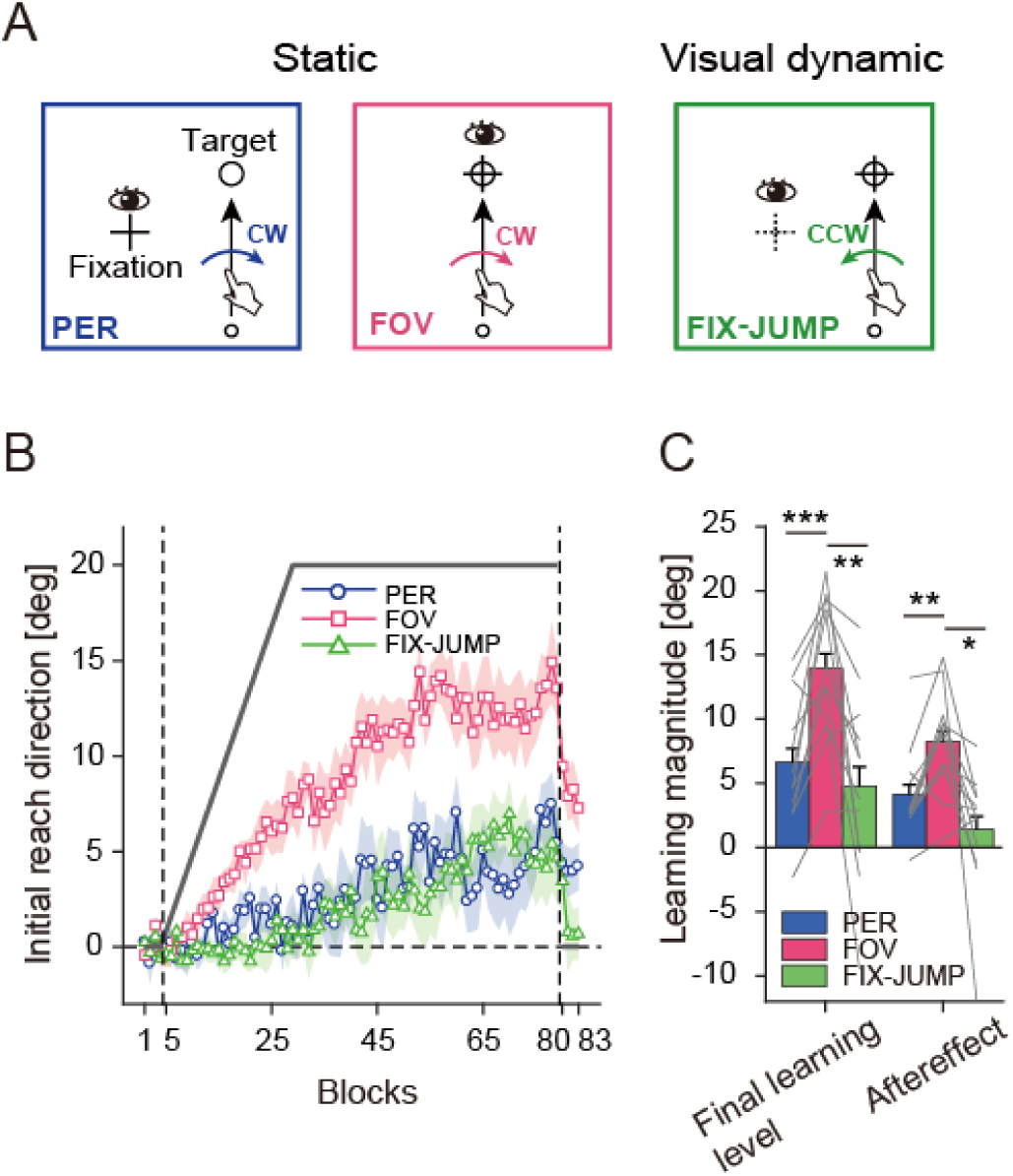
Control Experiment 1: Testing whether visual cues alone account for the simultaneous learning observed in Experiment 1. **A.** Trial conditions. PER and FOV were identical to those in Experiment 1. In FIX-JUMP, the fixation cross jumped to the target, but participants were required to maintain fixation at the original location and reach toward the target, resulting in the same gaze behavior as in PER. The direction of rotation was associated with static and visual dynamic conditions (i.e., CW for PER and FOV, CCW for FIX-JUMP). **B.** Learning curves under each condition (Initial reach direction; mean ± SE across participants). Figure formats are the same as in Figures 1D. **C.** Final learning level and aftereffect magnitudes. Bar graphs show the mean across participants (± SE), with gray lines indicating individual participant data. Asterisks indicate significant differences (Tukey post hoc tests; * p<0.05, ** p<0.01, *** p<0.001).

As in Experiment 1, PER, FOV, and FIX-JUMP were randomly presented within each block of 16 trials (4 trials each for PER and FOV and 8 trials for FIX-JUMP). Participants completed a pre-learning session (4 blocks/64 trials), learning session (75 blocks/1200 trials), and post-learning session (4 blocks/64 trials). In the pre- and post-learning sessions, normal visual feedback was presented. During the learning session, a visuomotor rotation was applied: the rotation angle gradually increased by 0.8° per block until it reached 20° then was maintained for the remainder of the session.

The spatial configurations of the reach target and fixation cross were the same as in Experiment 1. Under PER and FIX-JUMP, the fixation was placed either left or right of the target, counterbalanced across participants.

#### Experiment 2

Experiment 2 was designed to investigate whether two opposing visuomotor rotations were able to be learned when each was associated with a different sequential eye movement. Twelve participants (9 females; age 25.8 ± 8.7, mean ± SD) took part in this experiment. In each trial, participants performed one of two sequential saccade patterns, followed immediately by a reach toward the foveal target (Figure 3A). Participants began each trial by placing the cursor at the start point and directing their gaze to a fixation cross located 1 cm above the start point (F0, Figure 3A). After maintaining fixation for 0.7 s, the reach target appeared 10 cm above the start position. At 0.6 s after target presentation, the fixation jumped to 2 cm upward and 6 cm to the left or 2 cm upward and 6 cm to the right (F1). Participants were asked to make a saccade to this new fixation position as fast as possible. Once this eye movement was detected, the fixation cross jumped again to the reach-target location (F2). Thus, participants executed the first saccade to F1 then second saccade to F2 (i.e., reach target) and maintained fixation on the target until the subsequent reaching movement was completed. Depending on the direction of the first saccade, two saccade conditions were defined: Left-side and Right-side. A beep was presented 1 s after target appearance, signaling participants to initiate the reach. This timing was set so that reaching would begin immediately after the termination of sequential saccades. The actual time between saccade end and reach initiation was 221 ± 66 ms for Left-side saccade and 222 ± 65 ms for Right-side saccade (mean ± SD across participants).

**Figure 3.**
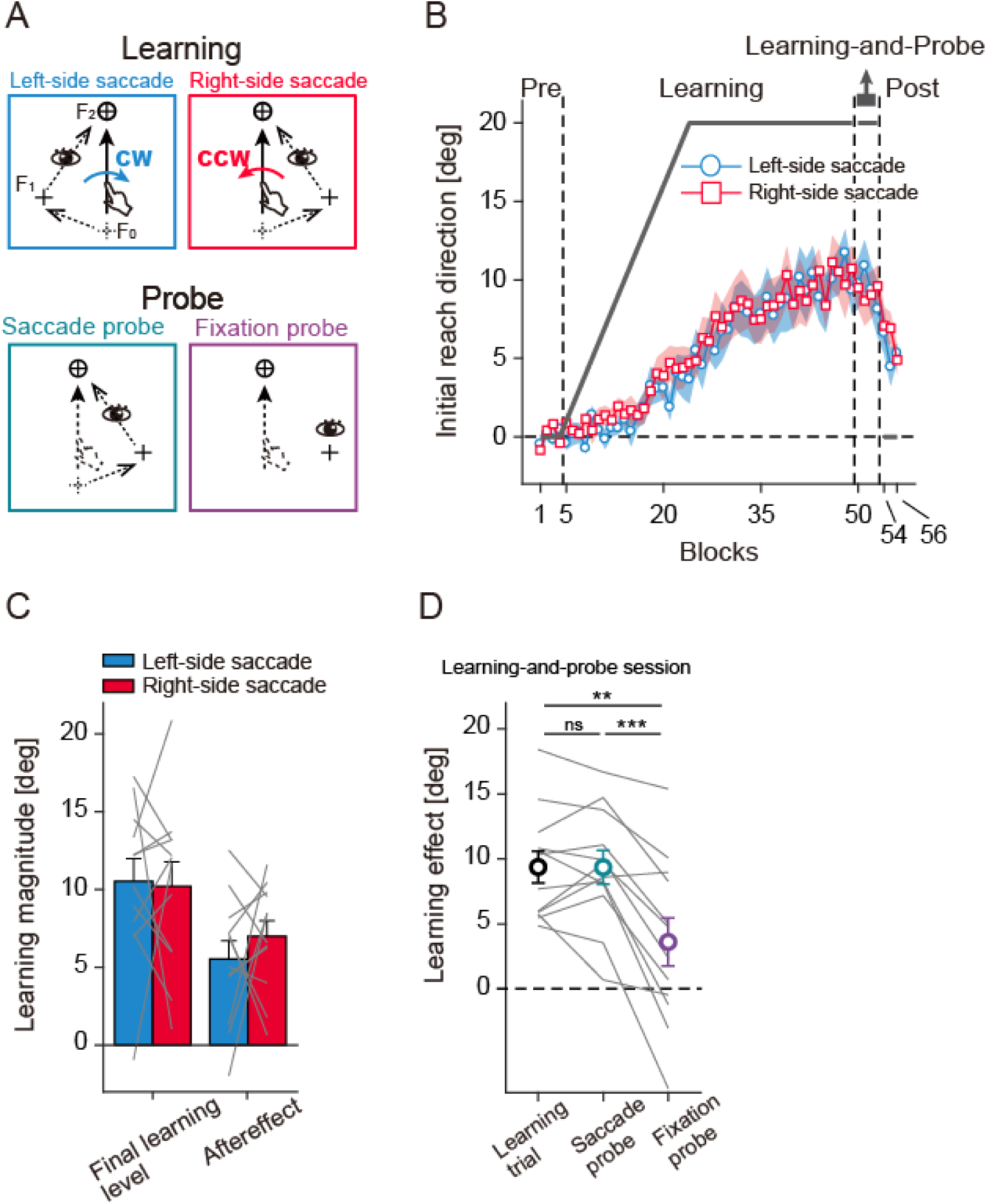
Experiment 2: Simultaneous learning based on different movement patterns of saccades. **A,** Trial conditions. In the learning session, participants performed either Left-side or Right-side sequential saccade (random order across trials), followed immediately by a reach toward the foveal target (upper panels). The direction of visuomotor rotation was associated with each condition. In the *learning and probe* sessions, two types of probe trials were additionally introduced (lower panels). In the *saccade probe*, participants executed the same eye-hand movements as in the learning trials, whereas in the *fixation probe*, they performed the reaching movement while maintaining fixation on an intermediate via point. In both probe trials, no cursor feedback was provided, enabling assessment of the retrieval of learned motor outputs under different gaze patterns. **B**–**C,** Learning curves for two saccade conditions (B) and final learning levels and aftereffect magnitudes (C). Data formats are the same as in Figures 1D and 1F, respectively. **D,** Learning effects in the learning and probe sessions. Initial reach direction averaged in this session was shown for learning trials, saccade probe, and fixation probe (gray lines: individual data; circles: group mean ± SEM). Asterisks denote statistically significant differences (** p<0.01 and *** p<0.001), and “ns” denotes non-significance.

In the learning session, the direction of visuomotor rotation was associated with the saccade conditions. For half the participants (n=6), CW was applied to the Left-side saccade, and CCW to Right-side saccade; this assignment was reversed for the other half. In the subsequent learning-and-probe session, two types of probe trials were interleaved with the learning trials to exclude the possibility that the static gaze state at F1, rather than dynamic sequential saccades, has a contextual effect on learning. The first, *saccade probe*, was identical to the learning trial, in which participants performed either Left-side or Right-side saccades, followed by reaching (Figure 3A). In the second, *fixation probe*, participants maintained gaze fixation at F1 before and during reaching movements. In both probe trials, no cursor feedback was provided. The type of eye movements (Left-side or Right-side saccade) in the saccade probe and the fixation location (left or right of the reach target) in the fixation probe were counterbalanced across participants.

The experiment was divided into four sessions. The first pre-learning session included two saccade conditions (Left-side saccade and Right-side saccade; normal cursor feedback) and two probe conditions (saccade probe and fixation probe; no cursor feedback). These four conditions were presented randomly within each block of 22 trials, consisting of 16 trials of the saccade conditions (8 trials for each) and 6 trials of the probe conditions (3 trials for each). The block was repeated four times (4 blocks/88 trials). In the subsequent learning session, two saccade conditions were presented randomly within each block of 16 trials (8 trials for Left-side saccade and 8 trials for Right-side saccade). The block was repeated 45 times (45 blocks/720 trials). As in Experiment 1, the angle of visuomotor rotation increased gradually by 0.8° per block up to 20° over the first 20 blocks and maintained 20° for the remaining 25 blocks. As mentioned above, the direction of rotation (CW or CCW) was uniquely associated with the saccade conditions. In the following learning-and-probe session, the block structure was identical to the pre-learning session: each block consisted of 22 trials, with 16 trials for the two saccade conditions (±20° rotation) and 6 trials for the two probe conditions (no-cursor feedback). The block was repeated three times (3 blocks/66 trials). The last post-learning session included only two saccade conditions with normal cursor feedback. Each block consisted of 16 trials (8 trials for each saccade condition) and was repeated 3 times (3 blocks/48 trials).

#### Control Experiment 2

Control Experiment 2 (twelve participants; 5 females; age 26.3 ± 6.1, mean ± SD) was conducted to rule out the possibility that visual cues from fixation jumps, rather than eye movements, served as contextual signals in Experiment 2. As in Experiment 2, each trial began when participants placed the cursor at the start point and fixated on the cross (F0; Figure 4A). After 0.6 s of target presentation, the fixation cross jumped to F1 and then to the reach target. Two conditions were defined based on the direction of the first jump: leftward and rightward displacement. The direction and amplitude of fixation jumps were identical to those in Experiment 2. Regardless of the direction of the fixation cross jump, participants were instructed to make a single saccade to the reach target as soon as the fixation cross appeared on it, followed immediately by a reaching movement. The actual time between saccade end and reach initiation was 154 ± 70 ms for Left-side and 152 ± 69 ms for Right-side conditions (mean ± SD across participants). The direction of visuomotor rotation was associated with the displacement condition (CW for leftward, CCW for rightward; shown in Fig. 4A), and this assignment was counterbalanced across participants. If visual cues from fixation jumps are sufficient for contextual learning, participants would exhibit simultaneous learning under these conditions.

**Figure 4.**
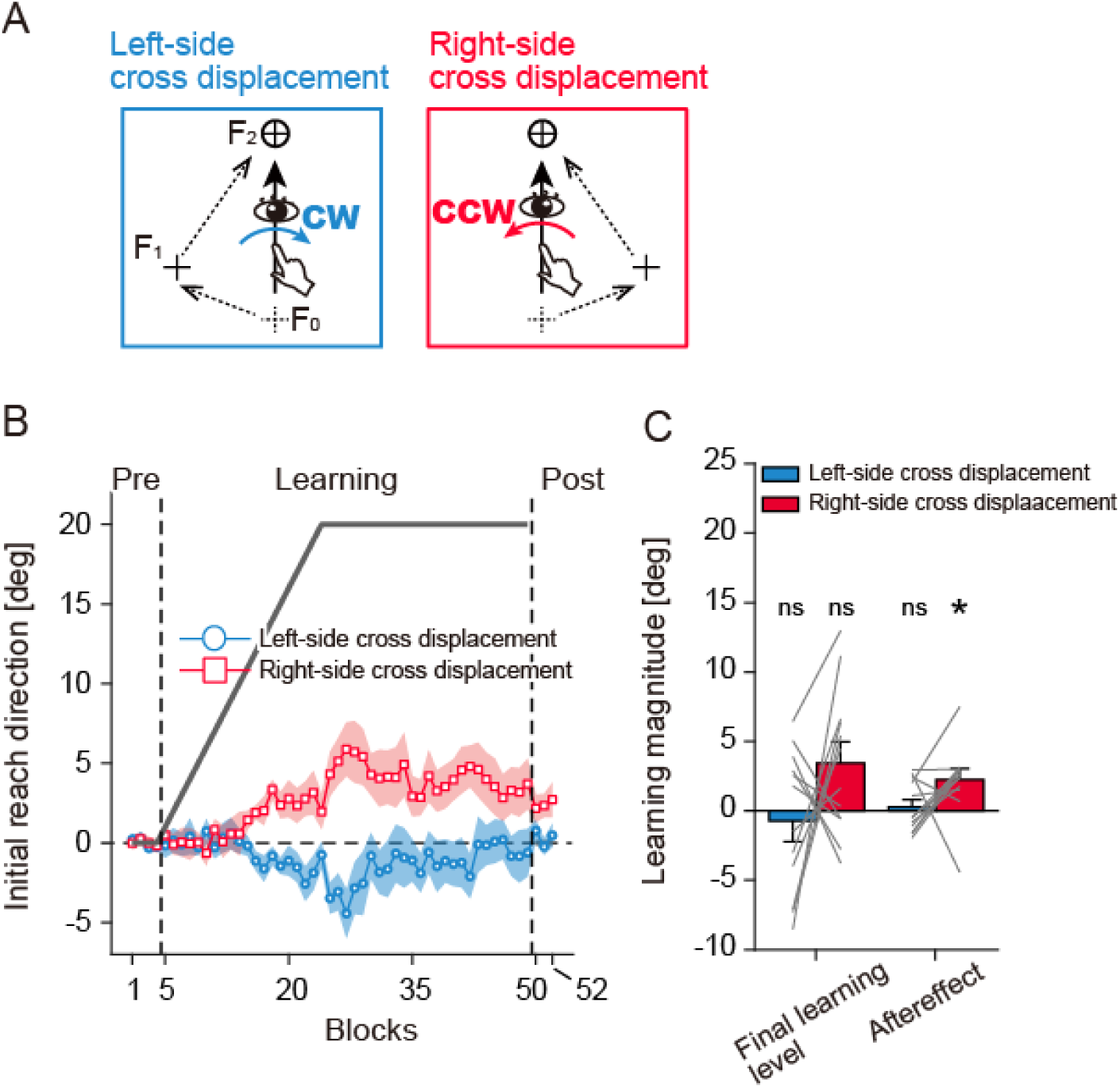
Control Experiment 2: Testing whether visual cues alone account for the simultaneous learning observed in Experiment 2. **A.** Task design and trial conditions. The movement of the fixation cross was identical to that in Experiment 2, and two conditions were defined by the direction of the first movement of the cross (left vs. right). Regardless of condition, participants were instructed to make a single saccade to the reach target upon fixation appearance, followed immediately by a reaching movement. The direction of rotation was associated with conditions (i.e., CW for left-side displacement and CCW for right-side displacement). **B.** Learning curves under each condition (Initial reach direction; mean ± SE across participants). Figure formats are the same as in Figures 1D or 3B. **C.** Final learning levels and aftereffect magnitudes. Bar graphs show the mean across participants (± SE), with gray lines indicating individual participant data. For both the learning level and the aftereffect, magnitudes were compared against zero using Bonferroni-corrected one-sample t-tests. Asterisks denote statistically significant differences (* p<0.05), and “ns” denotes non-significance.

Two conditions were randomly presented within each block of 16 trials (8 trials per condition). Participants completed a pre-learning session (4 blocks/64 trials), learning session (45 blocks/720 trials), and post-learning session (3 blocks/48 trials). In the pre- and post-learning sessions, normal visual feedback was provided. During the learning session, the rotation angle gradually increased by 1° per block until it reached 20°, after which it was maintained for the remainder of the session.

A gap duration of 0.18 s was inserted between the first and second fixation cross jumps. This interval was set to match the median interval between the first and second fixation jumps in Experiment 2, where the second jump was triggered by detection of the first saccade.

#### Experiment 3

Experiment 3 investigated the degree of learning transfer from a single training gaze state to other novel gaze states. Twelve participants (7 females; age 24.7 ± 8.7, mean ± SD) reached toward a target under 14 different gaze states (Figure 5B). Under the five static gaze conditions (blue 10–14 in Figure 5B), participants maintained the fixation throughout the trial, which was located at -15, -10, -5, 0, or 10 cm to the reach target along the horizontal direction. These fixation positions are indicated with blue dots in Figure 5B, with values in brackets. Under the nine dynamic gaze conditions (pink 1–9 in Figure 5B), participants made a horizontal saccade, followed immediately by reaching toward the target. In Figure 5B, the fixation positions before and after saccade are indicated with pink dots with their values.

**Figure 5.**
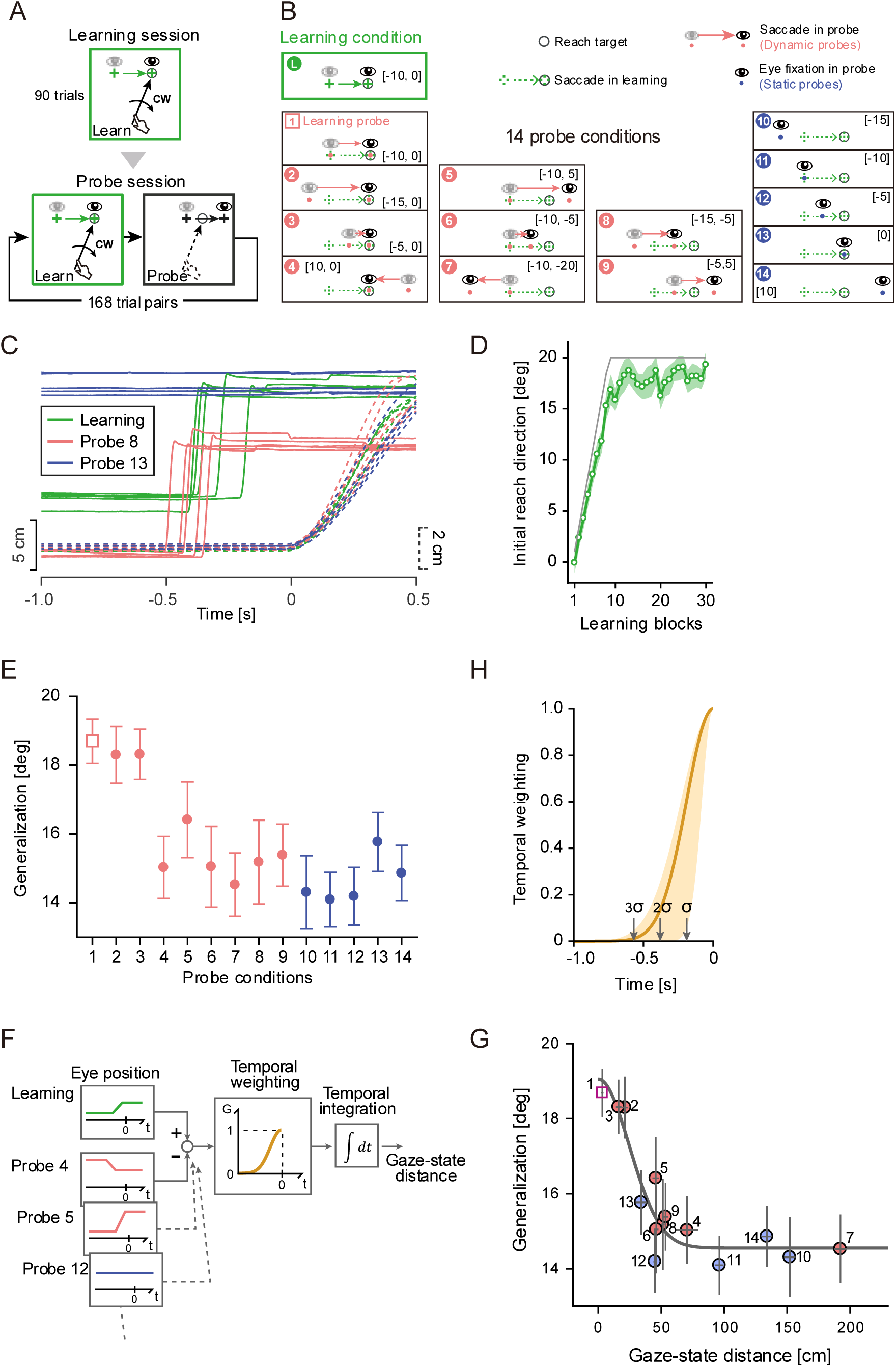
Experiment 3: Learning generalization across static and dynamic gaze states. **A,** Session design. In the learning session, participants made a saccade to the reach target, followed immediately by a reach toward the foveal target. They adapted to a visuomotor rotation (single direction; either CW or CCW). In the subsequent probe session, generalization of learning was tested across various gaze states, with no cursor feedback provided during reaching. To prevent memory decay, learning and probe trials were consistently presented in series. **B,** Fourteen gaze states tested in the probe session. Probe 1 was identical to the learning condition, in which reaching was preceded by a rightward saccade of 10 cm. Information about eye movements for dynamic probes (Probe 1–9) is denoted in brackets, indicating the horizontal start and end gaze positions relative to the reach target. For example, [-10, 0] in Probe 1 indicates that gaze was initially fixed 10 cm to the left of the reach target then shifted to the target location. Under the static probe conditions (Probes 10–14), values in brackets denote fixation location relative to the reach target. **C,** Example pattern of eye-hand movements. Solid lines show horizontal eye position (scale on the left), and dashed lines show forward hand position (scale on the right), both plotted relative to reach onset. **D,** Learning curve in the learning session. Initial reach direction (mean ± SE across participants) is plotted across blocks. The thin line indicates the angle of visuomotor rotation. **E,** Generalization in the probe session. Initial reach direction was measured in the probe trials without cursor feedback, reflecting retrieval of learned motor outputs. See panel B and the main text for details of the 14 probe conditions. Data are presented as mean ± SE across participants. **F,** Schematic for computing gaze-state distance. The gaze-state distance for each probe condition was calculated relative to the gaze state obtained in the learning trial. The difference in horizontal eye position (time-aligned to reach onset) was first computed and squared at each time point. These differences were then temporally weighted, with the weight decreasing as the time from reach onset increased. Finally, the temporally weighted differences were integrated over time. **G,** Generalization against gaze-state distance. Data for the 14 probe conditions (shown in panel B) were replotted as a function of gaze-state distance with a fitted curve. **H,** Estimated temporal weighting function (Gaussian). The function is shown with 95% confidence intervals based on bootstrap resampling. The arrows indicate three levels of the standard deviation of the Gaussian function. See main text for details.

The experiment began with a baseline session in which participants reached toward a single target located 10 cm above and 5 cm to the right of the start point, under 14 gaze conditions, as described above. Each block consisted of these 14 different gaze conditions, performed with and without cursor feedback, for a total of 28 trials in random order. The block was repeated 12 times (12 blocks/336 trials). In the feedback trials, veridical online cursor feedback was provided.

In the subsequent learning session, participants adapted to a single visuomotor rotation (CW or CCW, counterbalanced across participants), as shown in Figure 5A. In all trials, participants performed a rightward saccade of 10 cm toward the reach target, followed immediately by a reach toward the foveal target (learning condition in Figure 5B). Each block consisted of five trials. In most trials (3/5), the reach target was placed at the same position as in the baseline session, with the initial fixation point located 10 cm to the left of the reach target (learning condition in Figure 5B). To prevent participants from making habitual movements, in the remaining trials (2/5), both the reach target and initial fixation point were shifted 2.5 cm to the left or right of the original learning condition (i.e., saccade amplitude was maintained at 10 cm). These five trials were randomly presented within each block, and the block was repeated 30 times (30 blocks/150 trials). The angle of visuomotor rotation gradually increased by 0.5° per trial until reaching 20° then was maintained at 20° for the remainder of the learning session.

The last probe session consisted of a learning trial and probe trial (168 trials each; for a total of 336 trials), always in succession (Figure 5A). The learning trial was identical to the original learning condition in the learning session (i.e., 20° rotation; learning condition in Figure 5B) and was inserted before every probe trial to prevent rapid decay of adaptation. In the probe trials, participants performed reaching movements to the same target position used under the learning condition, under 14 gaze conditions (Figure 5B) without cursor feedback. These probe trials were identical to the non-cursor feedback trials in the baseline session.

### Statistical Analysis

The initiation and termination of reaching movements were detected online during the experiment and defined as the first time point at which tangential velocity exceeded or fell below 2 cm/s for at least 30 ms. For offline analysis, the obtained position data were aligned at the reach onset and low-pass filtered at 20 Hz (4th-order Butterworth, single sided). Velocity profiles were also obtained by three numerical time differentiations.

To assess learning, the initial direction of reaching movements (initial reach direction) was quantified on each trial as the angular difference between the movement trajectory and a line connecting the start point to the target. The initial reach direction was computed as the mean direction of velocity vectors during the 100–120-ms window after reach onset. To collapse data across different rotation directions (CW and CCW), the sign of the reach direction was adjusted so that positive values indicated compensatory movements for the corresponding rotation.

To remove movement biases across gaze conditions, we first averaged the initial reach direction under each condition in the pre-learning session. In Experiment 3, such biases were estimated across different gaze and visual conditions (cursor visible or invisible). These baseline biases were then subtracted from the corresponding trials throughout the experiment. This process was conducted for each participant.

The sign-adjusted reach direction, normalized by baseline, was used in subsequent analyses, including plotting learning curves and quantifying final learning levels, aftereffect magnitudes, and outputs of probe trials. In Experiment 1 and Control Experiment 1, the final learning level was defined as the mean reach direction in the last four blocks of the learning session, and the aftereffect magnitude as the mean reach direction in the four blocks of the post-learning session. In Experiment 2 and Control Experiment 2, these variables were estimated from the last two blocks of the learning session and from the first two blocks of the post-learning session. This difference in the number of blocks used for evaluation reflects differences in the number of trials per condition within each block across experiments, thereby ensuring a roughly comparable number of trials for evaluation.

Statistical analyses were performed in Jamovi 2.7.18. In Experiment 1, learning levels and aftereffects were analyzed using two-way mixed ANOVAs with Gaze Condition (PER, FOV, and SACCADE) as a within-participants factor and Gap Duration (Short, Middle, and Long) as a between-participants factor. Post hoc comparisons were performed using Tukey’s honestly significant difference (HSD) test. Details of the statistical design for Experiment 1 are provided in the Results section describing Figure 1E and G. To further confirm learning effects, learning levels and aftereffects in each gaze condition were tested against zero using Bonferroni-corrected one-sample t-tests (See the descriptions of Figure 1F and H).

For the analyses of Experiment 2 and Control Experiment 1, learning levels and aftereffects were analyzed using one-way repeated-measures ANOVAs followed by Tukey’s HSD tests. In Experiment 2, Control Experiment 1, and Control Experiment 2, learning levels and aftereffects were also tested against zero using Bonferroni-corrected one-sample t-tests.

We further conducted comparisons between the control experiments and their corresponding main experiments. Final learning levels were compared between Experiment 1 and Control Experiment 1 using a two-way mixed ANOVA, with gaze condition as a within-participants factor and experiment as a between-participants factor. The same analysis was performed to compare Experiment 2 and Control Experiment 2. Additional details specific to each comparison are provided in the corresponding Results sections.

For the analysis of Experiment 3, the generalization magnitude for each gaze probe was compared with that for the reference learning probe using Dunnett’s test (See the descriptions of Figure 5E).

## Results

### Experiment 1

In Experiment 1, as described in the experimental paradigm in the Method section, we investigated whether humans are able to learn two opposing internal models when each was associated with either a static or dynamic gaze condition. Participants performed reaching movements toward a single target under three gaze conditions: reaching to a peripheral target while maintaining fixation (PER), reaching to a foveated target under fixation (FOV), and reaching after a target-directed saccade (SACCADE; Figure 1B). During learning, CW and CCW visuomotor rotations were changed across trials, but were associated with either a static or dynamic gaze condition (i.e., CW for PER and FOV; CCW for SACCADE; Figure 1B). This association manner was counterbalanced across participants. Given that distinct static gaze conditions (i.e., PER and FOV) serve as effective contextual cues for simultaneous learning of opposing visuomotor rotations (Abekawa et al., 2022), we designed three gaze conditions (PER, FOV, and SACCADE) to isolate the contribution of eye movements, as explained in the Method section.

To assess learning, we quantified the initial direction of reaching movements. The sign of reach directions was adjusted for each gaze condition so that a positive value indicates the compensating direction against each visuomotor rotation (CW or CCW). As shown in Figure 1D (left), participants in the Short group exhibited clear learning progress under all three gaze conditions. In the post-learning session, reach directions remained positive despite the restoration of normal visual feedback, indicating clear aftereffects. Consistent with our prediction, in the Middle and Long groups (Figure 1D, center and right), learning under FOV and SACCADE (green and pink curves) appeared to be impaired, whereas learning under PER remained largely unchanged (See Experiment 1 in the Materials and Methods for details of the hypothesis regarding the temporal effect).

To quantify these observations, the final learning level was calculated from the last four blocks of the learning session. A two-way mixed ANOVA was conducted with Gaze condition (PER, FOV, and SACCADE; within participants) and Gap duration (Short, Middle, and Long; between-participants) as factors. As shown in Figure 1E, the analysis revealed significant main effects of Gaze condition (F(2,66)=14.78, p<0.001) and Gap duration (F(2,33)=3.63, p=0.038), as well as a significant interaction between Gaze condition and Gap duration (F(4,66)=8.71, p<0.001). Specifically, this interaction indicates that the effect of gap duration on learning differed across gaze conditions.

Returning to the results of the Short group (Figure 1D, left), learning progressed similarly across all three gaze conditions. The final learning level was significantly different from 0 under all conditions (Figure 1F, Short; PER: t(11)=9.5, p-adj.=3.5×10⁻⁶; FOV: t(11)=9.7, p-adj.=3.0 × 10⁻⁶, SACCADE: t(11)=15.7, p-adj.=2.1 × 10⁻⁸; Bonferroni-corrected t-tests) and did not differ significantly between gaze conditions (Figure 1E; simple main effect of Gaze: F(2,22)=0.037, p=0.96). These results indicate that static and dynamic gaze contexts can serve as contextual cues for separating the learning of conflicting visuomotor rotations.

For the Middle group (Figure 1D, center), learning still progressed under all conditions, as the final learning levels remained significantly greater than 0 (Figure 1F, Middle; PER: t(11)=9.9, p-adj.=2.5×10⁻⁶; FOV: t(11)=6.8, p-adj.=8.7×10⁻^5^, SACCADE: t(11)=7.1, p-adj.=5.7×10^-5^). However, the final learning levels differed significantly among gaze conditions (simple main effect of Gaze: F(2,22)=7.68, p=0.003). As shown in Figure 1E, Tukey post-hoc tests revealed that learning for PER was significantly greater than for FOV (t(11)=4.43, p=0.003), whereas no significant difference was observed between PER and SACCADE (t(11)=1.15, p=0.50) or between FOV and SACCADE (t(11)=−2.39, p=0.09). These results indicate that the inserted gap duration induced learning interference for FOV and SACCADE, leading to a partial failure of simultaneous learning.

The learning impairments in FOV and SACCADE observed in the Middle group were more pronounced in the Long group (Figure 1D, right). The final learning level differed significantly between gaze conditions (simple main effect of Gaze: F(2,22)=19.0, p<0.001). As shown in Figure 1E, Tukey post-hoc tests revealed that learning for PER was significantly greater than for both FOV (t(11)=5.89, p<0.001) and SACCADE (t(11)=5.32, p<0.001). In contrast, no significant difference was observed between FOV and SACCADE (t(11)=1.78, p=0.22). In addition, the final learning level was significantly different from zero for PER and FOV, but not for SACCADE (Figure 1E, Long; PER: t(11)=12.9, p-adj.=1.6×10^-7^; FOV: t(11)=5.1, p-adj.=0.001, SACCADE: t(11)=2.1, p-adj.=0.18). These results suggest that inserting a longer gap between saccade termination and reach initiation increased the overlap between the contextual gaze cues available in the SACCADE and FOV conditions, thereby enhancing interference between the two visuomotor maps and impairing learning.

We next examined how the inserted gap duration affected learning. As shown in Figure 1E, the final learning levels for FOV and SACCADE were reduced when a gap duration was introduced (simple main effect of Gap duration, FOV: F(2,33)=4.46, p=0.019, SACCADE: F(2,33)=8.94, p<0.001), whereas learning for PER was unaffected (F(2,33)=0.29, p=0.75). Post-hoc tests further revealed that learning in the Long group was reduced relative to the Short group for FOV (t(33)=2.8, p=0.02) and relative to both the Short (t(33)=3.8, p=0.002) and Middle (t(33)=3.5, p=0.004) groups for SACCADE. These results suggest that, as described above, inserting the gap duration reduced the distinctiveness of the FOV and SACCADE contexts, leading to learning interference. In contrast, PER remained a distinct contextual cue, allowing learning to proceed largely unaffected by the gap duration.

The patterns observed in the final learning level were generally reflected in the aftereffects estimated from the four blocks of the post-learning session (Figure 1G, H). A two-way mixed ANOVA revealed significant main effects of Gaze condition (F(2,66)=4.42, p=0.016) and Gap duration (F(2,33)=7.22, p=0.003), as well as a significant interaction (F(4,66)=4.21, p=0.004). Notably, in the Short group, aftereffect magnitudes significantly differed from 0 for all gaze conditions (p-adj.<2.0×10^-5^) and did not differ across conditions (simple main effect of gaze: F(2,22)=0.85, p=0.44). These results indicate successful context-dependent retrieval of visuomotor mappings corresponding to the immediately preceding gaze states. Consistent with the results for the final learning level, when a gap duration was inserted (i.e., in the Middle and Long groups), aftereffects for FOV and SACCADE were reduced, whereas those for PER remained largely unaffected. In particular, in the Long group, aftereffects differed between gaze conditions (simple main effect of gaze: F(2,22)=7.55, p=0.003; Figure 1G) and the aftereffect for SACCADE was not significantly different from 0 (t(11)=2.58, p-adj.=0.08; Fig.1 H).

Before drawing conclusions from Experiment 1, we considered the possibility that differences in reaching parameters across gaze conditions, rather than gaze behavior, might have served as learning contexts. To test this, we analyzed reaching kinematics during the pre-learning session in the Short group. However, no parameters significantly differed between SACCADE and PER and between SACCADE and FOV (Supplementary Table 1). These results rule out a reach kinematics-based explanation and suggest the contribution of distinct static and dynamic gaze contexts to reach learning. Notably, the contextual effect of dynamic gaze states decays over time.

### Control Experiment 1

We further conducted a control experiment to test if the visual cue of fixation jump, rather than eye movements, could facilitate learning. The paradigm was the same as in Experiment 1, except that SACCADE was replaced with the fixation jump condition (FIX-JUMP) (Figure 2A). Under FIX-JUMP, the fixation cross jumped to the target, but participants were instructed to maintain their gaze at the original fixation location without making any eye movements. As in Experiment 1, one direction of visuomotor rotation was associated with two static conditions (i.e., PER and FOV), and the other direction was associated with FIX-JUMP. If visual cues are sufficient for simultaneous learning, comparable learning should be observed across all three conditions, as in the Short group of Experiment 1. In contrast, if gaze states, rather than visual cues, are crucial, learning should fail for PER and FIX-jump, as these conditions share the same gaze states (i.e., reaching to a peripheral target) while being associated with opposing visuomotor rotations (CW and CCW; See Fig. 2A). Meanwhile, learning should progress for FOV, as it has a unique gaze state (i.e., reaching to a foveal target).

Consistent with latter prediction, the results showed that learning was evident only under FOV, whereas PER and FIX-JUMP showed minimal improvement, indicating a failure of simultaneous learning (Figure 2B). This pattern was statistically confirmed by quantifying the final learning level and aftereffects (Figure 2C). A one-way repeated-measures ANOVA revealed a significant effect of condition on final learning levels (F(2,22)=10.7, p<0.001). Post hoc Tukey tests confirmed that FOV was greater than both PER (t(11)=4.9, p=0.001) and FIX-JUMP (t(11)=3.9, p=0.006). Aftereffect magnitudes showed a similar pattern: a significant effect of condition (F(2,22)=6.8, p=0.005), with FOV greater than both PER (t(11)=4.6, p=0.002) and FIX-JUMP (t(11)=3.3, p=0.02).

We further compared the final learning levels between the Short group in Experiment 1 and Control Experiment 1 using a two-way mixed ANOVA, with gaze condition as a within-participants factor (PER, FOV, and SACCADE in Experiment 1; PER, FOV, and FIX-JUMP in Control Experiment 1) and experiment as a between-participants factor. Note that SACCADE in Experiment 1 corresponded to FIX-JUMP in Control Experiment 1. The analysis revealed significant main effects of experiment (F(1,22)=8.47, p=0.008), gaze condition (F(2,44)=7.84, p=0.001), and a significant Gaze x Experiment interaction (F(2,44)=8.44, p<0.001). Importantly, post hoc Tukey tests showed that learning was significantly smaller in PER (t(22)=3.20, p=0.042) and FIX-JUMP (t(22)=3.78, p=0.012) of Control Experiment 1 than in corresponding PER and SACCADE of Experiment 1, whereas no significant difference was found for FOV between experiments (t(22)=0.39, p=0.999). These results clearly indicate that visual cues alone were insufficient to facilitate simultaneous learning. Whereas PER and FIX-JUMP, which shared the same gaze state (i.e., reaching to peripheral target), exhibited little learning due to memory interference, FOV, which had a distinct gaze state (i.e., reaching to foveal target), showed robust learning. The results of Experiment 1 and Control Experiment 1 suggest that the distinction between static and dynamic gaze states, rather than visual cues, serves as a crucial contextual cue for separating competing motor memories.

### Experiment 2

Experiment 1 provided evidence that competing visuomotor maps can be separated when a dynamic gaze state is paired with static fixation states, suggesting that the distinction between static and dynamic gaze states can serve as a contextual cue for simultaneous learning. However, this design did not establish whether different dynamic gaze states themselves are sufficiently distinct to support separate motor memories. Experiment 2 was designed to address this question directly.

In each trial, participants performed one of two types of sequential saccades, followed immediately by reaching toward the target (Figure 3A, top). Each trial began with gaze fixation at F0, after which the fixation cross jumped to F1 then to F2. Participants were asked to make successive saccades from F0 to F1 then F1 to F2 as quickly and smoothly as possible. The final fixation point (F2) always coincided with the reach target. Therefore, participants performed reaching toward a foveal target while maintaining fixation on F2 throughout the reach.

In the learning session, the direction of visuomotor rotation was pseudo-randomly selected across trials, but uniquely associated with the saccade condition (i.e., CW for Left-side saccade and CCW for Right-side saccade; Figure 3A). The results indicate that participants gradually changed the reach direction over trials for both saccade conditions (Figure 3B). Final learning levels were significantly greater than 0 for both conditions (p-adj.<8.9×10^-5^) and did not differ between them (paired t-test, t(11)=0.16, p=0.88; Figure 3C). Similarly, aftereffect magnitudes were greater than 0 (p-adj.<0.0013 for both conditions) and did not differ between conditions (t(11)=0.89, p=0.39).

In this task, the initial and final fixation points (F0 and F2 in Figure 3A) were identical across the two saccade conditions, indicating that memory separation had to rely on different eye-movement patterns. An alternative possibility, however, was that the static gaze state at the intermediate point (F1) during the two sequential saccades might act as a contextual cue. To test this, we introduced a learning-and-probe session following the learning session, in which two learning and two probe conditions were interleaved across trials (Figure 3A). The learning conditions replicated the trials in the learning session (i.e., Left-side and Right-side saccade). The probe conditions included a saccade probe, in which participants executed sequential saccades followed by reaching, and a fixation probe, in which participants maintained fixation on F1 before and during reaching (Figure 3A, bottom). Under both probe conditions, no visual feedback of the cursor was provided, enabling assessment of memory retrieval. If learning relied on dynamic gaze states, reach directions comparable to learning trials would appear in the saccade probes; if it relied on static gaze states they would appear in the fixation probes.

The results indicate that reach directions for the saccade probe were comparable to those for the learning trial, whereas those for the fixation probe were smaller (Figure 3D). One-way ANOVA revealed a significant main effect of condition (learning trials, saccade probe, fixation probe; F(2,22)=18.7, p<0.001). Post hoc Tukey comparisons showed that the saccade probe was not different from learning trials (t(11)=0.03, p=1.0), whereas the fixation probe was different from both learning trials (t(11)=4.3, p=0.003) and the saccade probe (t(11)=5.2, p<0.001). These results indicate that distinct dynamic gaze states (i.e., different eye movement patterns) preceding reach initiation can support the separation of two opposing motor memories for reaching.

### Control Experiment 2

Regarding the results of Experiment 2, one possibility remains: learning may be facilitated by different visual patterns of fixation cross jumps. To test the contextual effect of such visual information, rather than eye movements, we conducted Control Experiment 2, in which a fixation cross jumped either to the left or right, as in Experiment 2 (Figure 4A). Regardless of the direction of the cross jump, participants were instructed to make a single saccade toward the target, immediately followed by reaching to the foveal target. The direction of visuomotor rotation was associated with the direction of cross movement.

If visual cues alone were sufficient to facilitate simultaneous learning, we would expect simultaneous learning in both conditions comparable to that observed in Experiment 2. As shown in Figure 4B, however, participants did not exhibit clear learning for either condition, likely because of memory interference. Crucially, the final learning level was not significantly different from 0 in either condition (Figure 4C; Left-side; t(11)=0.49, p-adj.=1, Right-side; t(11)=2.25, p-adj.=0.09). Regarding aftereffect magnitudes, the Right-side condition differed from 0 (t(11)=2.9, p-adj=0.03), whereas the Left-side condition did not (t(11)=0.50, p-adj=1.0), again indicating a failure of simultaneous learning.

We further compared the final learning levels between Experiment 2 and Control Experiment 2 using a two-way mixed ANOVA, with condition (Left-side and Right-side) as a within-participants factor and experiment as a between-participants factor. The analysis revealed a significant main effect of experiment (F(1,22)=43, p<0.001), whereas neither the main effect of condition (f(1,22)=1.38, p=0.25) nor the interaction (F(1,22)=1.88, p=0.184) was significant. The significant reduction in learning observed in Control Experiment 2 indicates that visual cues are substantially less effective as contextual cues for learning than gaze-state cues.

Consistent with Control Experiment 1, these findings again suggest that visual information associated with fixation-cross jumps is insufficient to support simultaneous learning. Rather, eye movements provide the contextual cue necessary for separating competing motor memories.

### Experiment 3

Finally, we investigated which factors in dynamic-gaze contexts contribute to reaching adaptation, using a generalization paradigm. Participants adapted to a single visuomotor rotation (20° CW or CCW; counterbalanced across participants). In each learning trial, they executed a rightward saccade toward the reach target, followed immediately by a reach to the foveal target (Figure 5A). In the subsequent probe session, we tested learning generalization to 14 possible gaze conditions (Figure 5B), with no cursor feedback. For the dynamic probes (probes 1–9 in Fig. 5B), reaching movements were preceded by saccades, the locations and amplitudes of which varied across probes. For the static probes (probes 10-14), reaches were performed while fixation was maintained at a certain location. Reach target remained fixed at the position used during the learning session. To prevent rapid decay of memory, each probe trial was preceded by a learning trial identical to those in the learning session (20° rotation; Figure 5A). For clarity, we plotted typical eye (horizontal position, solid lines) and hand (forward position, dashed lines) movements from the learning and selected probe conditions to illustrate the temporal eye-hand relationship (Figure 5C).

As in Experiments 1 and 2, we measured the sign-adjusted initial reach direction. A positive value indicated appropriate compensation for the rotation in the learning session and retrieval of learned movements in the probe session. As shown in Figure 5D, reach directions gradually increased over the course of the learning session, indicating clear progress in learning. In the subsequent probe session, the largest reach direction was observed for the learning probe, which had the same gaze state as under the learning condition (probe 1; pink square in Figure 5E). We regarded this learning probe as a reference and compared it with the remaining 13 probe conditions using Dunnett’s test. Large reach directions were found for dynamic probes 2 and 3, which did not differ significantly from the learning probe (p=0.99 for both; Figure 5E). These probes included saccades with the same endpoint and direction as in the learning condition (Figure 5B). Other dynamic gaze probes showed significantly lower reach directions, indicating limited transfer of learning. Specifically, these probes shared only endpoint (probe 4; p<0.001), start point (probes 5, 6, and 7; p<0.05, p<0.001, and p<0.001, respectively), or amplitude (probes 8 and 9; p<0.001 for both) with the learning condition. These results suggest that learning of reaching movements is robustly transferred between dynamic gaze conditions when the preceding saccade shares both direction and endpoint.

Experiment 1 suggested that dynamic and static gaze states may activate distinct neural states for separating motor memories. Consistent with this view, we found that learning transfer from the saccade-accompanied learning condition to static probes realized by fixation was limited (Figure 5B, probes 10–14). As shown in Figure 5E, the reach directions for these static probes were significantly smaller than for the learning probe (p<0.001 for all). Notably, under the learning condition, the eye fixated on the reach target after saccade, and its gaze state was identical as that for static probe 13 (see green and blue solid lines in Figure 5C). Thus, the limited transfer at probe 13 further supports the contribution of eye movements preceding each reach during reach learning.

In comparison between static probes, the reach direction for probe 13 (fixating the target) appeared greater (i.e., greater learning transfer) than for other static probes (probes 10, 11, 12, and 14). For the dynamic probes, as already noted, saccade endpoint and direction—rather than start point or amplitude—were crucial for learning transfer. Taken together, these results indicate that learning transfer depends on the recent history of gaze states before reach onset. To estimate such periods and capture the generalization pattern, we developed a model to compute gaze-state distance between the learning condition and each probe condition. As shown in Figure 5F, for each probe condition, the squared difference from the learning condition was calculated at each time point, using the horizontal eye position aligned to reach onset. The squared difference was temporally weighted by a Gaussian function with mean 0 and standard deviation *σ_t_*:

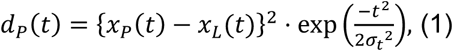

where *x_P_*(*t*) and *x_L_*(*t*) denote the horizontal eye position at time *t* for the probe *P* and learning condition, respectively. For temporal weighting, only the first half of the Gaussian was used, with the gain set to 1 at time 0 (i.e., reach onset) and decreased with earlier time points. The time-weighted distance of *d_P_*(*t*) was then integrated over time, and the square root of this integral was defined as the gaze-state distance of probe *P* (*D_P_*) from the learning condition, as follows:

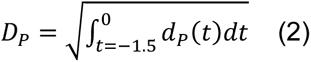

Due to the data-recording window, time integration was executed between -1.5 and 0 s. Figure 5G shows the reach direction plotted against the calculated gaze-state distance for each probe condition. The data were fitted with a Gaussian function as follows:

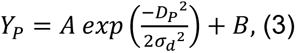

where *Y_P_* is the estimated initial reach direction for probe *P*, *A* is the gain factor, *B* is the overall offset, and *σ_d_* is the standard deviation of the Gaussian. The center of the Gaussian was fixed at 0, and the latter half of the function was used. In this model approach, four free parameters were estimated: the width of the temporal weighting (*σ_t_*) and three parameters for the generalization curve *(A*, *B*, and *σ_d_*), using a nonlinear least-squares fitting. The model fit to the group data is shown over the observed data in Figure 5G (coefficient of determination, R^2^ =0.87).

Our main interest in this model approach was to estimate the duration of preceding gaze states involved in the encoding of motor memories. Such duration is expressed as *σ_t_*, which was estimated at 0.19 s. Figure 5H shows the temporal weighting function with 95% confidential intervals based on bootstrap resampling (N = 2000). Considering a threshold of three times the standard deviation, our results indicate that the temporal history of gaze states until 0.57 s (confidence interval: 0.24–0.69 s) before reaching onset contributed to the learning of reaching movements. The fitting results and 95% confidence intervals for the four parameters were as follows: *A*=4.5 [3.5-5.6], *B*=14.6 [13.0-15.9], *σ_d_*=24.6 [2.4-32.3], and *σ_t_*=0.19 [0.08-0.23].

## Discussion

Reaching movements are typically preceded by coordinated eye movements. Such coordination has primarily been studied in relation to the roles of retinal and extraretinal signals in target representation and reaching accuracy (Prablanc et al., 1979; Henriques et al., 1998; Binsted and Elliott, 1999; Soechting et al., 2001; Buneo et al., 2002). Other studies have focused on reaction times of eye and hand movements to elucidate the interconnected mechanisms between both motor systems (Fisk and Goodale, 1985; Sailer et al., 2000; Dean et al., 2011; Armstrong et al., 2013; Yttri et al., 2013). Surprisingly little attention has been paid to how preceding eye movements impact reach-memory encoding, despite the prevalence of eye-hand coordination during motor learning in daily life. Using context-dependent and generalization learning paradigms, we demonstrated that preceding dynamic gaze states contribute to the formation and updating of internal models for reaching.

Several studies using sequential reaching tasks have reported improvement in reach learning when participants were allowed free eye movements rather than gaze fixation (Vieluf et al., 2015; Massing et al., 2016). However, these effects may reflect richer visual information, such as target location and error signals, obtained by eye movements (Gouirand et al., 2019). Thus, prior studies could not dissociate whether the effect was due to the inherent linkage of gaze states to the reach internal model or to the visual effect on the online reach control. Similarly, although we previously showed that different gaze conditions, reaching toward foveal and peripheral targets, can separate visuomotor memories (Abekawa et al., 2022), potential effects of differences in visual inputs during and/or at the end of the reaching movement could not be excluded. In the present study, gaze was directed at the reach target during reach execution, even under different gaze conditions associated with opposing visuomotor rotations (i.e., FOV and SACCADE in Experiment 1, Left-side and Right-side in Experiment 2). Thus, the observed simultaneous learning cannot be explained by the difference in the quality of online visual feedback control. Moreover, in Experiment 3, learning transfer remained limited in some probe conditions (e.g., probes 4 and 13), despite gaze being directed to the target during movement execution as in the learning condition (Fig. 5E). These results indicate that the preceding dynamic gaze states, rather than differences in visual feedback, play a significant role in the formation and expression of reaching memories.

The contextual encoding of multiple motor memories is a central issue in motor neuroscience (Heald et al., 2021, 2023). Arbitrary visual cues generally fail to facilitate learning (Gandolfo et al., 1996; Howard et al., 2013), as confirmed in our Control Experiments (but, see Refs. 27 and 28). In contrast, effective contextual cues are typically related to the state of the hand (Gandolfo et al., 1996; Ghahramani and Wolpert, 1997; Nozaki et al., 2006; Hirashima and Nozaki, 2012; Howard et al., 2013, 2015) or hand-associated objects (Hwang et al., 2003; Heald et al., 2018; Proud et al., 2019). Previous work (Nozaki et al., 2006) demonstrated that, in a bimanual reaching task, force-field learning in one arm was impacted by concurrent movements of the other arm but not by ankle flexion, suggesting that contextual states should be somatotopically close to the trained effector. However, our findings, along with our previous study (Abekawa et al., 2022), show that gaze states—despite being somatotopically distant from the arm—can contribute to reach learning. These results challenge the importance of somatotopic proximity and instead highlight functional relevance to reach planning as the critical determinant of context-dependent reach learning. Gaze locations relative to the reach target provide essential information for target representation, and dynamic gaze states update such gaze-centered representations in reach planning (Henriques et al., 1998; Batista et al., 1999; Medendorp et al., 2003; Khan et al., 2005), which might contribute to learning. The importance of functional relevance is also supported by recent findings that decision uncertainty can serve as a learning context (Ogasa et al., 2024).

The impact of prior movements on motor learning suggests that adaptation depends not only on the current body states but also on its recent history (Wainscott et al., 2005; Howard et al., 2012, 2024; Gippert et al., 2023). This suggests that the learning effect is sensitive to the time interval between the contextual and adaptation movements, rather than being determined solely by the presence or absence of eye movements. Consistent with this view, the contextual learning effect in Experiment 1 (Figure 1 E and F) decreased from the Short (∼260-ms interval) to the Middle group (∼540 ms) and disappeared in the Long group (∼1200 ms). In Experiment 3, the generalization pattern was well accounted for a temporally weighted measure of gaze-state distance (Fig. 5G), indicating that learning incorporated gaze history up to approximately 600 ms before reach onset (Fig. 5H). This temporal window was broadly consistent with Experiment 1.

Several explanations may account for this temporal decay. One possibility is that the limited time window reflects a common mechanism across different learning paradigms. In sequential reaching tasks, the contextual effect of lead-in movements decays as the interval between contextual and adapted movements increased, disappearing by ∼600 ms (Wainscott et al., 2005; Howard et al., 2012). Similarly, associative learning between preceding contextual stimuli (sound or light) and reaching movements occurs at stimulus–reach intervals of a few hundred milliseconds but not beyond 1000 ms (Avraham et al., 2022). These findings, together with ours, suggest that prior contextual information influences memory encoding for subsequent reaching within a constrained temporal window. One potential mechanism underlying this temporal window is cerebellar processing, as frequently investigated in eye-blink conditioning, a well-established cerebellum-dependent learning paradigm that is highly sensitive to the timing between a conditioned (e.g., tone or light as context) and unconditioned stimulus (e.g., air puff eliciting a blink response). Consistent with previous motor learning studies and our findings, optimal conditioning occurs when the conditioned stimulus precedes the unconditioned stimulus by approximately 500 ms (e.g., Ref. (Kjell et al., 2018)).

Alternatively, the temporal window in gaze-dependent learning may stem from eye-movement-specific processes. To separate motor memories by preceding gaze states, the brain needs to retain saccade-related information for a certain period to influence reach planning and/or execution. Neurophysiological studies have shown that the dorsolateral prefrontal cortex and lateral intraparietal cortex (LIP), both involved in saccade control, exhibit post-saccadic activity lasting approximately 500 ms after saccade termination (Goldman-Rakic et al., 1990; Funahashi et al., 1991; Takeda and Funahashi, 2002; Genovesio et al., 2007; Funahashi, 2014; Graf and Andersen, 2014). Within the first 300 ms, this post-saccadic activity carries information about the visual location of saccade target, saccade direction (Funahashi et al., 1991; Takeda and Funahashi, 2002), and movement vector in depth (Genovesio et al., 2007). Eye position and saccade direction were also accurately decoded by population activities in the LIP after saccade termination (Graf and Andersen, 2014). Thus, post-saccadic activity could provide contextual signals for reaching memories, with its impact decaying over time after the saccade. Consistent with our findings, a recent behavioral study suggested that preceding saccades are used as learning contexts in a sequential saccades task (Azadi and McPeek, 2022). Together, these results raise an interesting question of whether the temporal features of contextual learning reflect gaze-specific processes or more general motor-learning mechanisms.

Several behavioral studies (Wainscott et al., 2005; Howard et al., 2012, 2024; Sheahan et al., 2016; Gippert et al., 2023), along with the present study, highlight the importance of contextual cues during reach planning for motor learning. These findings align with the framework that views motor cortices as a dynamical system (Shenoy et al., 2013). Preparatory neural activities in the premotor and primary motor cortices influence reach learning (Vyas et al., 2020; Sun et al., 2022). Human imaging (Ogawa and Imamizu, 2013) and brain stimulation (Nozaki et al., 2016) studies further supported the critical role of motor cortices in context-dependent learning. Accordingly, our results suggest that preceding gaze states induce distinct preparatory neural states for reaching, enabling the formation of multiple motor memories. Consistent with this interpretation, gaze-dependent neural signals have been identified along the dorsal visuomotor pathway, including premotor areas involved in eye-hand coordination (Boussaoud et al., 1998; Jouffrais and Boussaoud, 1999; Fujii et al., 2000; Cisek and Kalaska, 2002; Pesaran et al., 2006, 2010; Batista et al., 2007). Future work is needed to determine how parietal, motor, and cerebellar circuits contribute to contextual encoding and motor-memory formation.

In daily situations such as playing tennis, individuals often learn and switch different reaching skills (e.g., forehand and backhand shots) while tracking a moving ball with their gaze. Given the limitations of external cues in contextual learning, preceding eye movements can serve as contextual cue for reaching, potentially facilitating motor-skill acquisition and expression. Considering the highly variable eye-hand coordination across behaviors, gaze-independent learning would of course be more appropriate than a gaze-dependent manner in certain situations. Important questions remain as to how the brain transitions between gaze-dependent and gaze-independent learning and how these mechanisms can be effectively applied in real-life settings, including sports and rehabilitation.

## ACKNOWLEDGMENTS

We thank T. Isobe for the data collection and A. Kimura, and N. Miyazaki for their support and encouragement.

---

**Supplementary Table 1.**
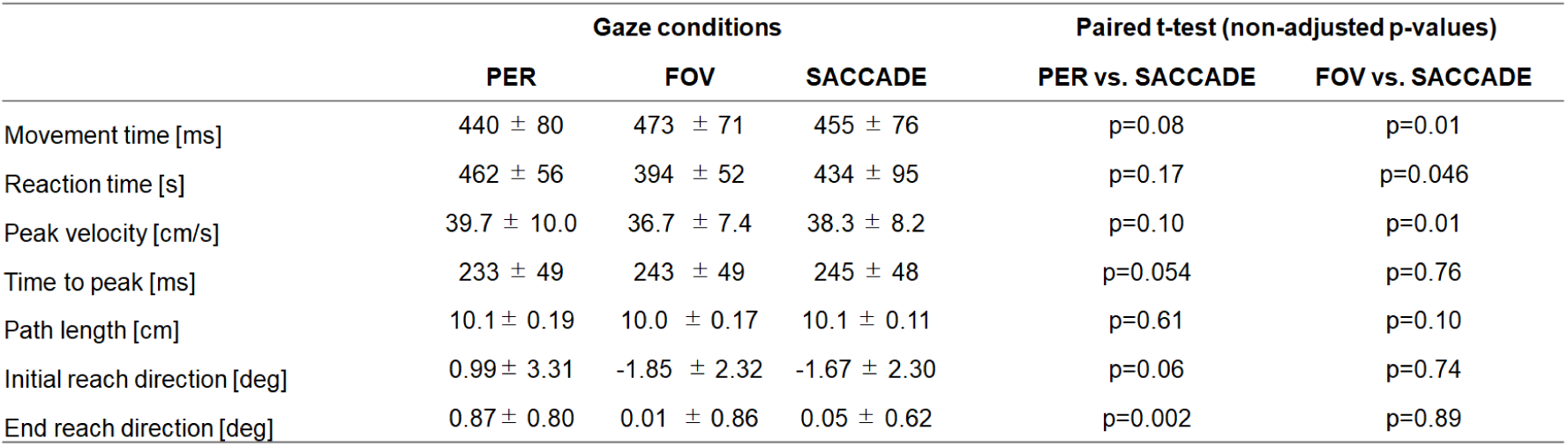
Reaching kinematics for short group during pre-learning session (normal visual feedback) in Experiment 1. Each movement-related parameter was compared between PER and SACCADE and between FOV and SACCADE using paired T-tests. A positive reach-direction value indicates a clockwise deviation from a straight path to the target. For each participant, measures were averaged across the four blocks of the pre-learning session. The table shows the mean ± SD across participants. No measure differed significantly in both the PRE-SACCADE and the FOV-SACCADE comparisons. Thus, the simultaneous learning observed under the three gaze conditions in Experiment 1 cannot be explained by a contextual effect of different reach-related parameters across conditions.

