## Supplemental Table 1 for "Eye-hand coordination beyond vision: preceding gaze history shapes reaching memories"

| | Gaze conditions | | | Paired t-test (non-adjusted p-values) | |
|---|---|---|---|---|---|
| | PER | FOV | SACCADE | PER vs. SACCADE | FOV vs. SACCADE |
| Movement time [ms] | 440 ± 80 | 473 ± 71 | 455 ± 76 | p=0.08 | p=0.01 |
| Reaction time [s] | 462 ± 56 | 394 ± 52 | 434 ± 95 | p=0.17 | p=0.046 |
| Peak velocity [cm/s] | 39.7 ± 10.0 | 36.7 ± 7.4 | 38.3 ± 8.2 | p=0.10 | p=0.01 |
| Time to peak [ms] | 233 ± 49 | 243 ± 49 | 245 ± 48 | p=0.054 | p=0.76 |
| Path length [cm] | 10.1 ± 0.19 | 10.0 ± 0.17 | 10.1 ± 0.11 | p=0.61 | p=0.10 |
| Initial reach direction [deg] | 0.99 ± 3.31 | -1.85 ± 2.32 | -1.67 ± 2.30 | p=0.06 | p=0.74 |
| End reach direction [deg] | 0.87 ± 0.80 | 0.01 ± 0.86 | 0.05 ± 0.62 | p=0.002 | p=0.89 |
